# Beyond modularity: Fine-scale mechanisms and rules for brain network reconfiguration

**DOI:** 10.1101/097691

**Authors:** Ankit N. Khambhati, Marcelo G. Mattar, Danielle S. Bassett

## Abstract

The human brain is in constant flux, as distinct areas engage in transient communication to support basic behaviors as well as complex cognition. The collection of interactions between cortical and subcortical areas forms a functional brain network whose topology evolves with time. Despite the nontrivial dynamics that are germaine to this networked system, experimental evidence demonstrates that functional interactions organize into putative brain systems that facilitate different facets of cognitive computation. We hypothesize that such dynamic functional networks are organized around a set of rules that constrain their spatial architecture – which brain regions may functionally interact – and their temporal architecture – how these interactions fluctuate over time. To objectively uncover these organizing principles, we apply an unsupervised machine learning approach called nonnegative matrix factorization to time-evolving, resting state functional networks in 20 healthy subjects. This machine-learning approach automatically clusters temporally co-varying functional interactions into subgraphs that represent putative topological modes of dynamic functional architecture. We find that subgraphs are stratified based on both the underlying modular organization and the topographical distance of their strongest interactions: while many subgraphs are largely contained within modules, others span between modules and are expressed differently over time. The relationship between dynamic subgraphs and modular architecture is further highlighted by the ability of time-varying subgraph expression to explain inter-individual differences in module reorganization. Collectively, these results point to the critical role subgraphs play in constraining the topography and topology of functional brain networks. More broadly, this machine learning approach opens a new door for understanding the architecture of dynamic functional networks during both task and rest states, and for probing alterations of that architecture in disease.

## 1. Introduction

More than just a sum of its parts, the brain performs computations and processes information by linking functionally specialized areas through complex patterns of anatomical wiring [2, 43, 120, 114]. Indeed, the underlying structural network forms the foundation of a wide repertoire of functional interactions between different regions [40, 107, 39]. Collectively, these interactions can be modeled as edges between nodes in a graph [5, 99, 25, 24, 62, 111] to probe the neurophysiological underpinnings of thought, perception, and action [112, 80]. Importantly, to actuate behavior and cognition through a changing landscape of environmental demands, these patterns of functional interactions must flexibly reconfigure [56, 26, 70, 77], presumably according to organizing principles that coordinate the dynamic engagement and disengagement of distinct sets of brain areas [9, 42, 89, 29].

A fundamental core of this dynamic architecture is thought to be *modularity* – the division of functionally engaged brain regions into putative modules that may compartmentalize computation within discrete functional systems – such as motor, visual, auditory, or attention – without disturbing brain regions in other systems [86, 113]. To robustly support functional dynamics driving behavior and cognition, functional brain networks capably reorganize their module composition across different time scales by integrating and segregating brain regions within and across brain systems [16, 77]. Moreover, functional brain networks exhibit flexibility in their module composition as they adapt to cognitive demands associated with completing a task [20, 22, 77, 115], processing linguistic stimuli [42, 29], or learning a new skill [9, 12, 13]. Notably, individual differences in flexibility are correlated with individual differences in learning [9, 48], working memory performance [22], and cognitive flexibility [22], which is particularly interesting in light of its role as an intermediate phenotype in schizophrenia [20].

Yet, while flexibility appears to be an important attribute of functional brain networks, a fundamental understanding of how network reconfiguration occurs, and what rules constrain the types of reconfiguration that characterize neural systems, is lacking. Do brain regions spontaneously arrange themselves into efficient modular configurations, or do functional interactions obey a distinct set of rules that constrain which modules can and cannot exist? Are there separable groups of functional interactions that differentially drive integration and segregation among network modules? While modules partition groups of brain regions based purely on the presence or strength of functional interactions, they remain agnostic to the *topology* of the functional interactions within and between them. These open questions have motivated the development of a new wave of graph theoretic tools grounded in machine learning that recover latent structure in dynamic brain networks as coherent groups of temporally co-varying functional interactions – known as subgraphs.

Mathematically, functional subgraphs form a basis set of unique patterns of graph edges whose weighted linear combination – given by a set of time-varying basis weights for each subgraph – reconstructs a repertoire of graph configurations observed over time. All graph nodes participate to varying degree in each subgraph, and the edges between nodes are assigned weights based on how strongly they co-vary over time. Thus, subgraphs are computationally represented as weighted adjacency matrices equal in size to the original graph – enabling us to query subgraph architecture as we might with brain graphs. More conceptually, functional subgraphs can be thought of as topological modes of interacting brain regions that are differentially expressed over time [72, 27]. Spatially, individual brain regions may engage and play distinct topological roles among different groups of brain regions in multiple subgraphs – a capability that is critical for capturing network architecture of putative functional brain systems [93, 27].

The framework of functional subgraphs yields an opportunity to examine subgraph topology and subgraph dynamics associated with different behavioral and cognitive states. For example, functional brain networks could decompose into different components of visual processing in which visual areas interact with ventral attention areas in one subgraph and with dorsal attention areas in another – with each subgraph increasing or decreasing its relative expression during different phases of cognitive processing. A recent application of the subgraph decomposition technique in neurodevelopment demonstrated that functional brain networks of children and young adults are composed of subgraphs representing known brain systems that are common across both age groups yet differ in the temporal properties of their expression during the resting state [27]. Such decomposition has also been used to uncover putative network subregions in epilepsy that play unique roles in the initiation and maintenance of neural dysfunction [63]. Despite the apparent role of subgraphs as functional substrates of information processing in the brain, their topographical and topological properties are not well understood. Do subgraphs reveal network architecture that is spatially distributed across the brain? Are subgraphs bound to the modular architecture of brain networks or do they span between modules?

Here, we develop a quantitative framework for identifying functional subgraphs and characterizing their relationship to whole brain network architecture. We base our analysis on a burgeoning application of non-negative matrix factorization (NMF) that decomposes dynamic functional networks into constituent additive parts rather than generalized features [71]. In implementing NMF in the context of neuroimaging data, we detail several important methodological considerations. Our flexible approach enables subgraph analysis across multiple spatiotemporal scales of network topology and dynamics through manipulation of three main parameters. By optimizing these parameters, we uncover a robust set of subgraphs across multiple human subjects and examine their sensitivity to functional interactions within and between network modules, and over different geographic distances. First, we hypothesize that subgraphs are selectively sensitive to functional interactions over different distances – short-range interactions are more likely to exist in some subgraphs and long-range interactions are more likely to exist in other subgraphs. This distance-wise stratification would underlie fundamental organization of brain networks into local, function-specific interactions (characteristic of clusters and modules [101, 116]) and distributed, integrative interactions (characteristic of hubs and rich-clubs [106, 38, 118]). Second, we expect that subgraphs are differentially sensitive to functional interactions within modules and functional interactions between modules. Functional interactions within the same module may undergo more similar patterns of temporal variation – and are therefore more likely to be clustered into one set of subgraphs – than functional interactions that span between modules – which are therefore more likely to cluster into another set of subgraphs. Subgraphs sensitive to network topology within modules might also exhibit a strong correlation between fluctuation in their temporal expression and flexibility of module reorganization over time. Such a relationship would highlight a novel perspective on the inter-regional changes of functional interactions that accompany meso-scale alterations in functional brain networks in both health and disease [20, 100, 44].

## 2. Methods

### 2.1. Experimental Design

Twenty participants (nine female; ages 19-53 years; mean age = 26.7 years) with normal or corrected vision and no history of neurological disease or psychiatric disorders were recruited for this experiment, non-overlapping results from which have been reported elsewhere [3, 79, 78, 53]. All participants volunteered and provided informed consent in writing in accordance with the guidelines of the Institutional Review Board of the University of Pennsylvania (IRB #801929).

### 2.2. Data acquisition and pre-processing

Magnetic resonance images were obtained at the Hospital of the University of Pennsylvania using a 3.0 T Siemens Trio MRI scanner equipped with a 32-channel head coil. T1-weighted structural images of the whole brain were acquired on the first of four scan sessions per subject using a three-dimensional magnetization-prepared rapid acquisition gradient echo pulse sequence (repetition time (TR) 1620 ms; echo time (TE) 3.09 ms; inversion time 950 ms; voxel size 1 mm × 1 mm × 1 mm; matrix size 190 × 263 × 165). A field map was also acquired at each of the four scan sessions (TR 1200 ms; TE1 4.06 ms; TE2 6.52 ms; flip angle 60°; voxel size 3.4 mm × 3.4 mm × 4.0 mm; field of view 220 mm; matrix size 64 × 64 × 52) to correct geometric distortion caused by magnetic field inhomogeneity. In all scans, T2*-weighted images sensitive to blood oxygenation level-dependent contrasts were acquired using a slice accelerated multiband echo planar pulse sequence (TR 500 ms; TE 30 ms; flip angle 30°; voxel size 3.0 mm × 3.0 mm × 3.0 mm; field of view 192 mm; matrix size 64 × 64 × 48).

### fMRI Preprocessing

We preprocessed the resting state fMRI data using FEAT (FMRI Expert Analysis Tool) Version 6.00, part of FSL (FMRIB’s Software Library, www.fmrib.ox.ac.uk/fsl). Specifically, we applied: EPI distortion correction using FUGUE [57]; motion correction using MCFLIRT [58]; slice-timing correction using Fourier-space timeseries phase-shifting; non-brain removal using BET [110]; grand-mean intensity normalization of the entire 4D dataset by a single multiplicative factor; highpass temporal filtering (Gaussian-weighted least-squares straight line fitting, with sigma=50.0s).

Nuisance timeseries were voxelwise regressed from the preprocessed data. Nuisance regressors included (i) three translation (X, Y, Z) and three rotation (Pitch, Yaw, Roll) timeseries derived by retrospective head motion correction (R = [*X, Y, Z, pitch, yaw, roll*]), together with expansion terms ([*RR*^2^*R*_*t*−1_R^2^_*t*−1_]), for a total of 24 motion regressors [46]); (ii) the five first principal components calculated from time-series derived from regions of non-interest (white matter and cerebrospinal fluid), using the anatomical CompCor method (aCompCor) [14] and (iii) the average signal derived from white matter voxels located within a 15mm radius from each voxel, following the ANATICOR method [60]. Global signal was not regressed out of voxel time series [85, 103, 30]. Finally, the mean functional image and the Harvard-Oxford atlas were co-registered using Statistical Parametric Mapping software (SPM12; Wellcome Department of Imaging Neuroscience, www.fil.ion.ucl.ac.uk/spm) in order to extract regional mean time-series (Fig. 1A).

**Figure 1:**
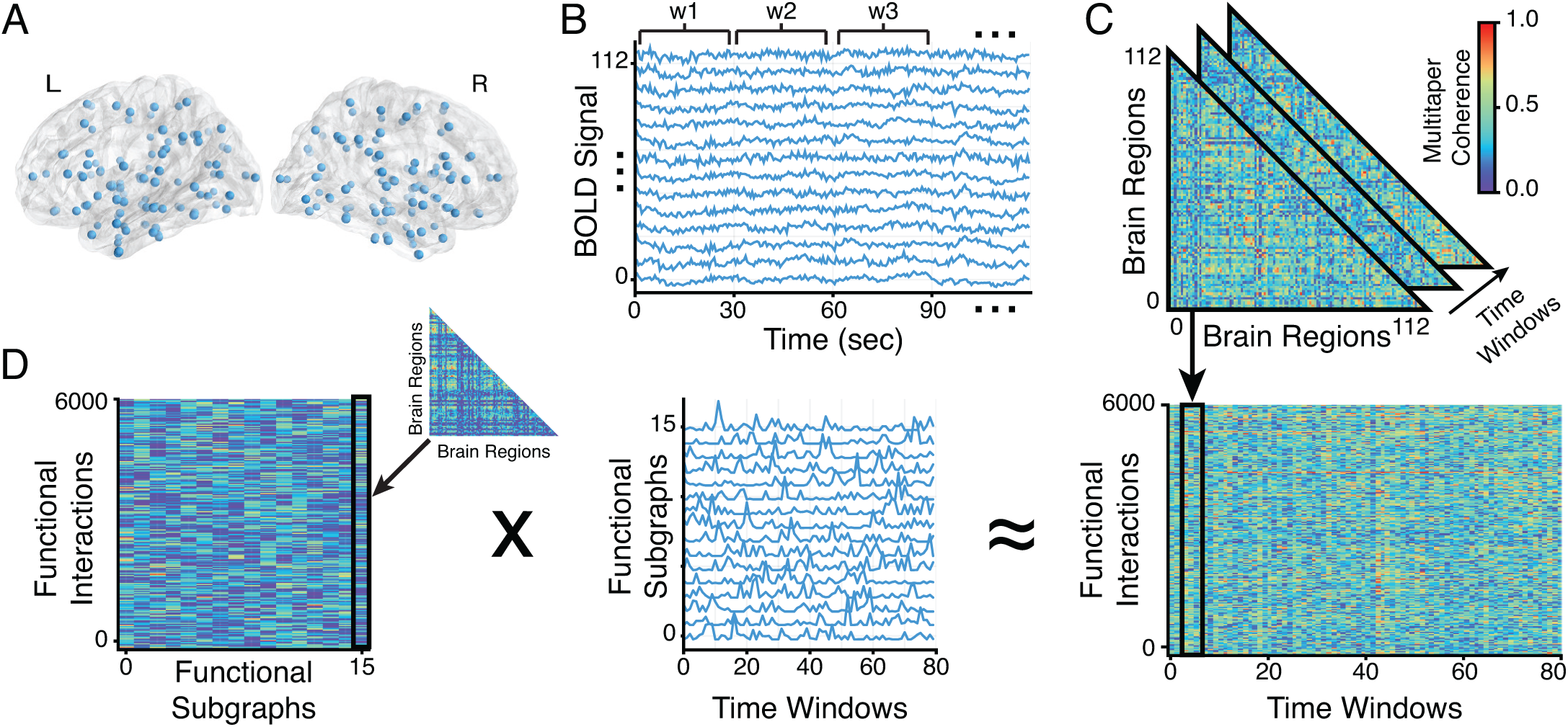
Subgraphs of dynamic functional brain networks. (*A*) We obtained multi-band fMRI BOLD signals from 112 functional regions of interest (HO-112 atlas) in cortical and subcortical areas during the resting state from 20 healthy subjects across four separate imaging sessions. We constructed dynamic functional networks for each subject by (*B*) dividing the regional BOLD signal into 80 non-overlapping time windows, each 30 seconds (60 TRs) in duration, and (*C*) computing multitaper coherence between each pair of regional BOLD signals for every time window to obtain a time-varying adjacency matrix – with brain regions as network nodes and time-varying coherence as weighted network edges. (*D*) We concatenated all pairwise edges over time and generated a time-varying network configuration matrix (*right*). We applied non-negative matrix factorization (NMF) – which pursues a parts-based decomposition of the dynamic network – to the configuration matrix and clustered temporally co-varying network edges into a matrix of subgraphs (*left*) and a matrix of time-dependent coefficients (*middle*) that quantifies the level of expression in each time window for each subgraph.

## 2.3. Constructing time-varying functional networks

To construct dynamic functional brain networks, we begin by dividing the BOLD signal over the four scan sessions into 80 non-overlapping time windows – each 30 seconds in duration and containing spectral information between frequencies of 0.03−1.00 Hz (Fig. 1B). For each time window, we measure functional interactions between each pair of brain regions based on coherence within a frequency band of 0.03−0.20 Hz. We use the *mtspec* Python implementation of multi-taper coherence [94] with time-bandwidth product of 2.5 and 4 tapers to achieve a frequency resolution of 0.08 Hz. Coherence values are stored in a time-varying, *N × N × T* adjacency matrix A, where *N =* 112 brain regions and *T =* 80 time windows, for each subject (Fig. 1C). In our weighted network analysis, we retain and analyze *all* functional interactions between brain regions, and do not apply any threshold or perform binarization.

An alternate representation of the three-dimensional network adjacency matrix **A** is a two-dimensional network configuration matrix **Â**, which tabulates all *N* × *N* pairwise interactions across *T* time windows. Due to symmetry of **A*_t_***, we unravel the upper triangle of **A*_t_***, resulting in the weights of 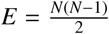 functional interactions. Thus, **Â** has dimensions *E* × *T*. We construct a separate network configuration matrix for each subject.

## 2.4. Partitioning the dynamic functional network

### 2.4.1. Clustering nodes into modules

To identify functional modules – partitions of highly inter-connected brain regions – we maximize the following multi-layer modularity quality function **𝙌** that assigns brain regions into modules that vary with time [84]:

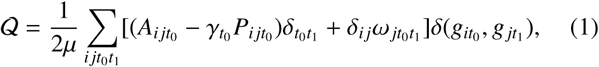

where indices *i*, *j* represent brain regions and *t*_o_, *t*_1_ represent consecutive time windows, *μ* is the sum of all functional interactions in the dynamic network, **P** represents functional interactions derived from a null model (e.g., the Newman-Girvan null model [87]), *γ* is a structural resolution parameter for modular organization within a single time window, *ω* is a temporal resolution parameter to model changes in modular organization over time, and *g* is the module assignment of a brain region within a time window [8]. In accord with prior studies, we choose resolution parameters such that *γ*_*t*0_ = *γ* = 1 and *ω*_*jt*_0_*t*_1__ = *ω* = 1 [9, 12, 21].

We use a Louvain-like greedy optimization to maximize **𝙌** [59]. The optimization landscape of the multi-layer modularity contains a plateau with many near-optimal solutions [49]. This near-degeneracy can be addressed by aggregating solutions over several runs of the Louvain algorithm [12]. To reach a consensus on module assignments, we separately apply 100 independent optimizations of **𝙌** to the dynamic functional network of each subject.

### 2.4.2 Clustering edges into subgraphs

To identify functional subgraphs – clusters of temporally co-varying network interactions — we apply an unsupervised machine learning algorithm called non-negative matrix factorization (NMF) [71] to the network configuration matrix (Fig. 1D). NMF decomposes the network configuration matrix into constituent subgraphs and accompanying time-varying expression coefficients [28, 63]. Each subgraph is an additive component of the original network – weighted by its associated time-varying expression coefficient – and represents a pattern of functional interactions between brain regions. The NMF-based subgraph learning paradigm is a basis decomposition of a collection of dynamic graphs that separates co-varying network edges into subgraphs – or basis functions – with associated temporal coefficients – or basis weights. Unlike other graph clustering approaches that seek a hard partition of nodes and edges into clusters [84, 8], the temporal coefficients provide a soft partition of the network edges, such that the original functional network of any time window can be reconstructed through a linear combination of all of the subgraphs weighted by their associated temporal coefficient in that time window [72, 73, 28, 63]. This implies that at a specific time window, subgraphs with a high temporal coefficient contribute their pattern of functional connections more than subgraphs with a low temporal coefficient.

Mathematically, NMF approximates **Â** by two non-negative matrices **W** – the subgraph matrix identifying patterns of functional interactions (with dimensions *E* × *m*) – and **H** – the time-varying expression coefficients matrix (with dimensions *m* × *T*) – such that:

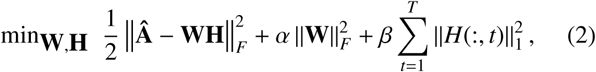

where *m* ∊ [2, min(*E*, *T*) – 1] is the number of subgraphs to decompose, *β* is a penalty weight to impose sparse temporal expression coefficients, and *α* is a regularization of the interaction strengths for subgraphs [66]. To solve the NMF equation, we use an alternating non-negative least squares with block-pivoting method with 100 iterations for fast and efficient factorization of large matrices [65]. We initialized **W** and **H** with non-negative weights drawn from a uniform random distribution on the interval [0,1].

To select the parameters *m*, *β*, and *α*, we pursue a random sampling scheme – shown to be effective in optimizing highdimensional parameter spaces [15] – in which we re-run the NMF algorithm for 30,000 parameter sets in which *m* is drawn from 𝓤(2,20), *β* is drawn from 𝓤(0.01,2), and *α* is drawn from 𝓤(0.01,2) (Fig. 2). We evaluate subgraph learning performance based on the following three criteria: residual error 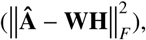 temporal sparsity 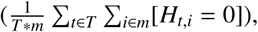 and subgraph sparsity 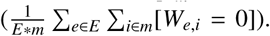 The optimal parameter set should yield subgraphs that (i) minimize the residual error and reliably span the temporal space of observed network topologies, (ii) maximize the temporal sparsity and effectively capture network architecture specific to distinct temporal states, and (iii) maximize the subgraph sparsity and pinpoint the strongest interactions between brain regions. Based on these criteria, we identified an optimum parameter set (*m̅*, *β̅*, *α̅*) that exhibits a low residual error in the bottom 25^*th*^ percentile, high temporal sparsity in the upper 25^*th*^ percentile, and high subgraph sparsity in the upper 25^*th*^ percentile of our random sampling scheme.

**Figure 2:**
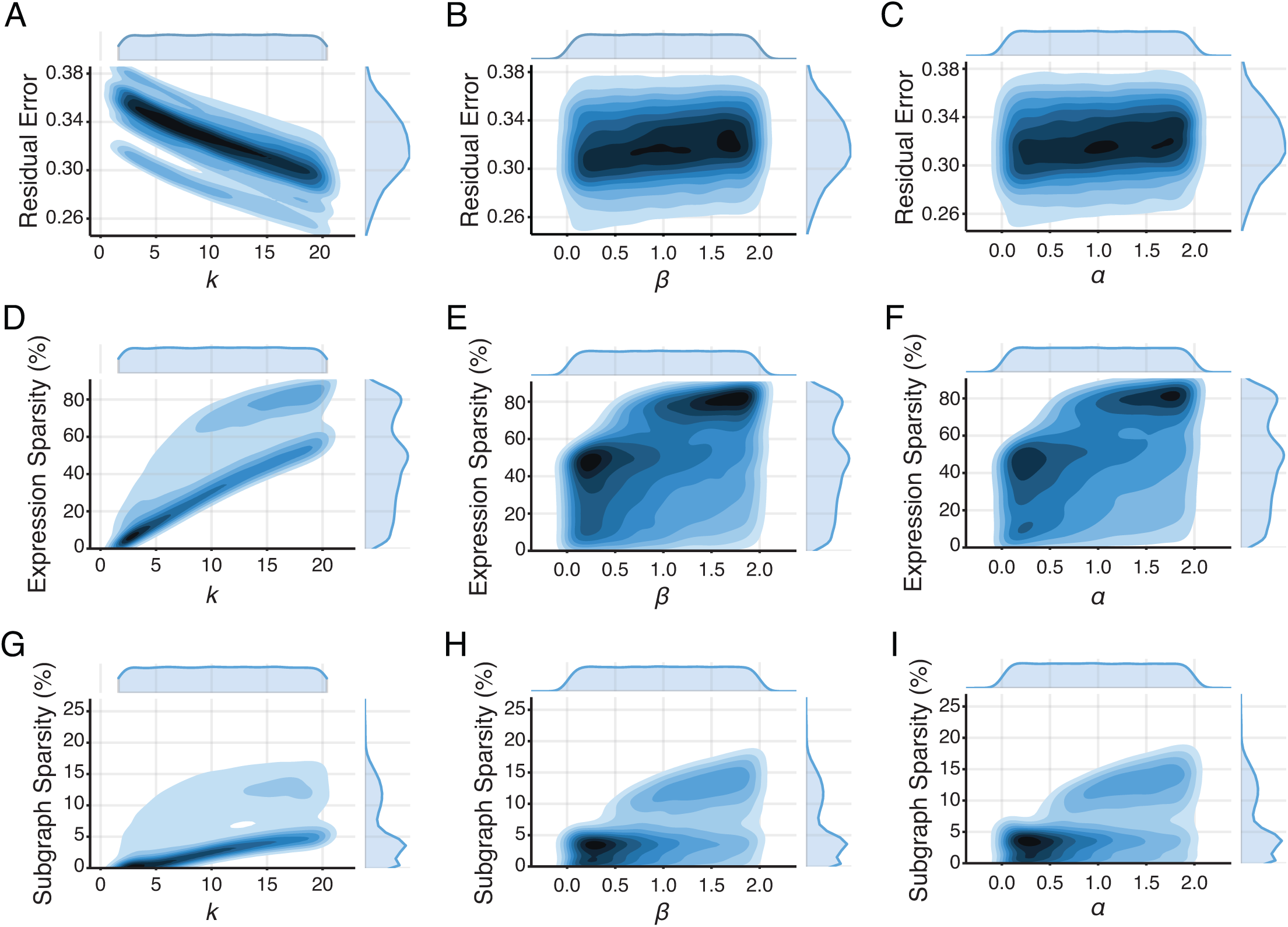
Parameter optimization for subgraph detection. NMF-based subgraph detection requires optimizing three parameters: the number of subgraphs *k*, the temporal sparsity of subgraph expression *β*, and the regularization of subgraph edge weights *α*. To characterize this parameter space, we randomly sampled *k*, *β*, and *α* from a three-dimensional uniform distribution (*k* ∊ [2,20], *β* ∊ [0.01,2.0], *α* ∊ [0.01,2.0]) and applied NMF to each subject using each parameter set. (*A-C*) The joint probability distribution of residual reconstruction error and marginalized *k* (Pearson *r* = −0.68, *p* < 1 × 10^−16^), marginalized *β* (Pearson *r* = 0.10, *p* < 1 × 10^−16^), and marginalized *α* (Pearson *r* = 0.11, *p* < 1 × 10^−16^) suggests that (i) increasing the number of subgraphs significantly improves the reconstruction of the original dynamic network, and (ii) increasing the subgraph expression sparsity or edge weight regularization slightly improves the reconstruction. (*D-F*) The joint probability distribution of percent sparse expression coefficients and marginalized *k* (Pearson *r* = 0.78, *p* < 1 × 10^−16^), marginalized *β* (Pearson *r* = 0.38, *p* < 1 × 10^−16^), and marginalized *α* (Pearson *r* = 0.37, *p* < 1 × 10^−16^) suggests that (i) increasing the number of subgraphs significantly introduces sparse temporal expression coefficients, and that (ii) increasing the subgraph expression sparsity of edge weight regularization moderately increases temporal expression sparsity. We observed two likely basins of low and high temporal expression sparsity for a range of *β* and a, centered around (*β* ∊ [0.1, 0.3], *α* ∊ [0.1,0.5]) and (*β* ∊ [1.25,2.0], *α* ∊ [1.25, 2.0]), suggesting that subgraphs may be specific to two different scales of temporal dynamics – local or global. (*G-I*) The joint probability distribution of percent sparse subgraph edge weights and marginalized *k* (Pearson *r* = 0.5, *p* < 1 × 10^−16^), marginalized *β* (Pearson *r* = 0.51, *p* < 1 × 10^−16^), and marginalized *α* (Pearson *r* = 0.51, *p* < 1 × 10^−16^) suggests that increasing the number of subgraphs, subgraph expression sparsity, and edge weight regularization increases the sparsity of subgraph edge weights.

Due to the non-deterministic nature of this approach, we integrated subgraph estimates over multiple runs of the algorithm using *consensus clustering* – a general method of testing robustness and stability of clusters over many runs of one or more non-deterministic clustering algorithms [83]. Our adapted consensus clustering procedure [52, 51] entailed the following steps: (i) run the NMF algorithm *R* times per network configuration matrix, (ii) concatenate subgraph matrix **W** across *R* runs into an aggregate matrix with dimensions *E* × (*R* * *m̅*), and (iii) apply NMF to the aggregate matrix to determine a final set of subgraphs and expression coefficients.

### 2.5. Measures of Subgraph Topology and Dynamics

To assess whether the functional interactions of a subgraph are constrained by network topography, we compute the Pearson correlation coefficient between the strength of functional interactions and the Euclidean distance between their associated brain regions, for each subgraph of each subject (Fig. 3). Positive correlations imply stronger functional interactions over longer distances and negative correlations imply stronger functional interactions over shorter distances. We also compute a surrogate distribution of Pearson correlation coefficient values for each subgraph by randomly permuting functional interactions between brain regions 1000 times.

**Figure 3:**
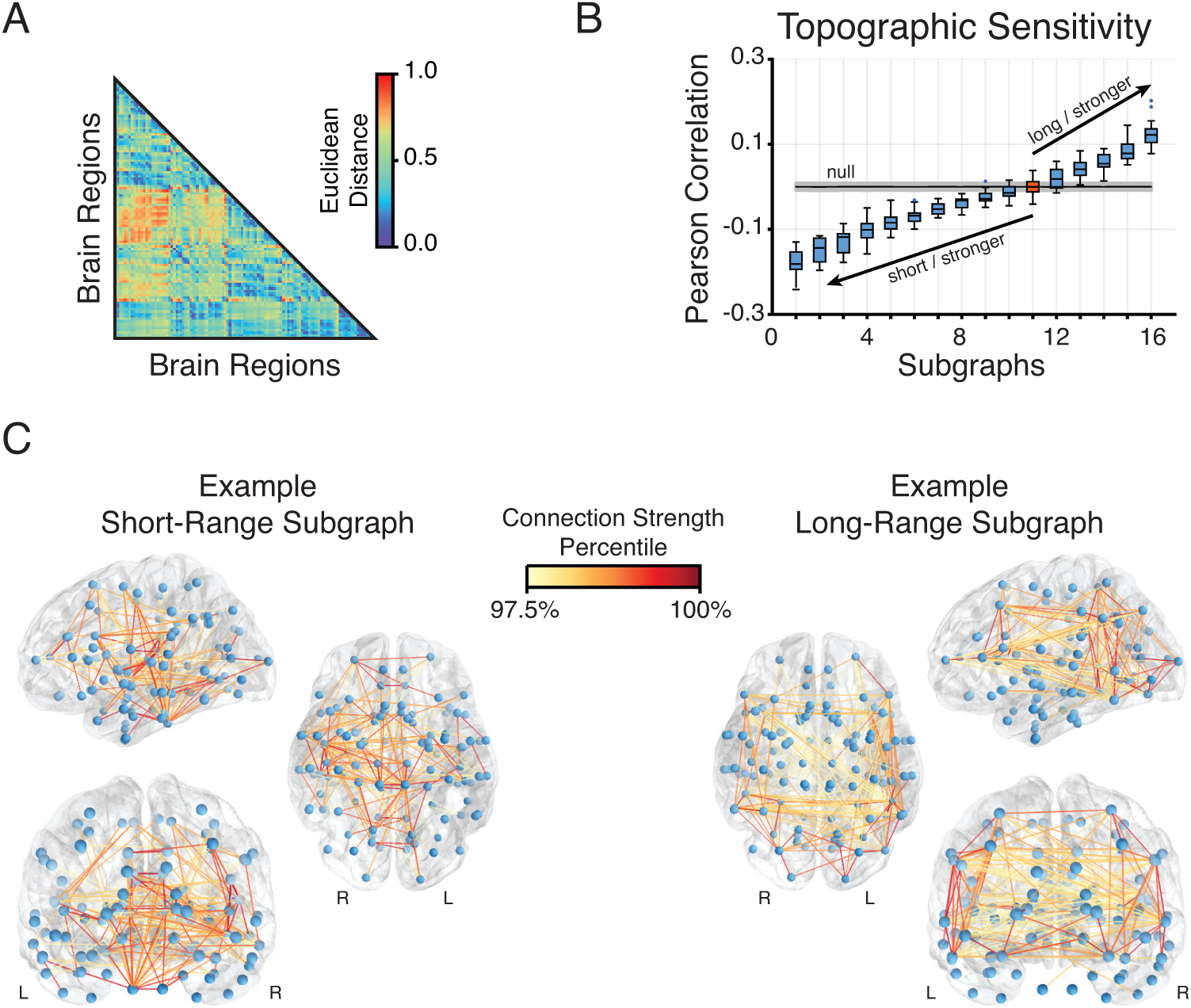
Topographical properties of subgraphs. (*A*) Topographic map derived from the Harvard-Oxford anatomical atlas composed of 112 areas in cortex, subcortex, and brainstem, representing the Euclidean distance between pairs of brain regions in physical space. (*B*) Distribution of topographical sensitivity for each subgraph – rank ordered from smallest to largest – across subjects. Topographical sensitivity is defined as the Pearson correlation coefficient between the topographic distance and subgraph-specific edge strength between all pairs of brain regions (see Methods): a more positive correlation indicates that edge strength increases with increasing distance between brain regions, whereas a more negative correlation indicates that edge strength increases with decreasing distance between brain regions. To identify subgraphs with significant topographical sensitivity, we generated a null model for each subgraph by randomly rewiring edge strengths between nodes and recomputing topographical sensitivity using a Pearson correlation. Compared to the null model, subgraphs 1-10 (blue) exhibited significantly negative topographical sensitivity, subgraphs 12-16 (blue) exhibited significantly positive topographical sensitivity and subgraph 11 (red) exhibited non-significant topographical sensitivity (Bonferroni corrected *t*-tests; *p* < 0.05). These results suggest that edge co-variances have a robust topographical organization in which the strongest edges may form subgraphs over either short or long geographical distances. (*C*) An example of a short-range subgraph with negative topographical sensitivity (*left*), and an example of a long-range subgraph with positive topographical sensitivity (*right*). The short-range subgraph exemplifies strong edges between nearby brain regions, whereas the long-range subgraph exemplifies strong edges between distant brain regions.

To determine whether the functional interactions of a subgraph are expressed within modules or between modules, we compute a time-varying module sensitivity index that maps dynamic module architecture onto subgraphs. Mathemati-cally, the module sensitivity index is computed as MSI(*m*, *t*) = 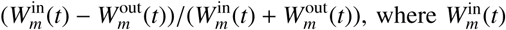 is the aver-age strength of functional interactions of the *m*^th^ subgraph between brain regions assigned to the same module at time *t*, and 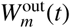 is the average strength of functional interactions between brain regions assigned to different modules at time *t*. We also compute a surrogate distribution of the module sensitivity index for each subgraph by permuting module assignments uniformly at random 100 times.

We compare subgraph dynamics to module reorganization by calculating the energy and skew of subgraph expression coefficients and the flexibility of module composition over time. The subgraph expression energy [28, 63] quantifies the overall magnitude expression of the subgraph and is based on the equation energy(*m*) = 𝔼[*H*_*m*_^2^], where *H_m_* are the temporal coefficients of the *m*^th^ subgraph. The skew of a distribution of subgraph expression coefficients quantifies how transiently or persistently subgraphs are expressed [28, 63]. Intuitively, transient subgraphs are expressed in brief, infrequent bursts – resulting in a heavy-tailed distribution of temporal coefficients (i.e., more small coefficients, and few large coefficients) – and persistent subgraphs are expressed evenly in time – resulting in a more normal distribution of temporal coefficients that fluctuate about the mean. The skew of the distribution of temporal coefficients for a subgraph distinguishes whether it is transiently (skew is greater than zero) or persistently (skew less than zero) expressed. The skew of the subgraph expression coefficients is computed based on the equation skew 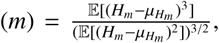 where *H_m_* are the temporal coefficients of the *m*^th^ subgraph and *μ_H_* is the mean of the coefficients. The module flexibility [10] is quantified by the mean fraction of times that brain regions change module assignment over the 80 time windows.

## 3. Results

### 3.1. Selecting parameters for NMF-based subgraph decomposition

Non-negative matrix factorization (NMF) requires parameter optimization that is critical for identifying a robust set of subgraphs that (i) best reconstruct the original dynamic network, and (ii) reflect functional interactions that are gradually expressed and not overly specific to a single point in time. We apply a random sampling scheme to characterize the rich parameter space of the number of subgraphs m, the temporal sparsity of subgraph expression *β*, and the regularization of subgraph edge weights *α* across the dynamic functional networks of 20 healthy subjects (Fig. 2).

First, we measure the relationship between residual reconstruction error and each parameter *m*, *β*, *α*, marginalizing over the remaining parameters (Fig. 2A-C). We observe a significant decrease in residual error as the number of subgraphs increases (Pearson *r* = −0.68, *p* < 1 × 10^−16^), suggesting that subgraphs collectively explain more of the statistical space of dynamic functional networks as the number of decomposed subgraphs increases. In contrast, we did not observe as strong of a relationship between residual error and the temporal sparsity parameter (Pearson *r* = 0.10, *p* < 1 × 10^−16^), or between residual error and the subgraph regularization parameter (Pearson *r* = 0.11, *p* < 1 × 10^−16^). Collectively, these results suggest that residual error is primarily driven by the number of subgraphs.

Importantly, we note a potential degeneracy associated with increasing the dimensionality of the subgraph sub-space *m* towards the dimensionality of the original network. For large *m*, we expect that subgraphs would sacrifice functional architecture that generalizes over time for architecture that is specific to single edges or single time windows. To characterize these degeneracies, we measure the percentages of sparse temporal expression coefficients and sparse subgraph edge weights as the number of subgraphs increases (Fig. 2D,G). We find a significant positive relationship between the percent of sparse temporal coefficients and the number of subgraphs (Pearson *r* = 0.78, *p* < 1 × 10^−16^), suggesting that for a large number of subgraphs, the temporal expression of subgraphs generally falls to zero and is compensated by brief increases in expression only at specific time points. Similarly, we find a significant positive relationship between the percent of sparse subgraph edge weights and the number of subgraphs (Pearson *r* = 0.5, *p* < 1 × 10^−16^), suggesting that for a large number of subgraphs, the subgraph topology approaches a degeneracy in which edge weights fall to zero with only a few non-zero edges constituting a subgraph.

To devise a strategy that would accommodate this degeneracy, we characterize the effect of *β* and *α* on the percent of sparse temporal coefficients (Fig. 2E,F) and the percent of sparse subgraph edge weights (Fig. 2H,I). We find moderate relationships between the percent of sparse temporal coefficients and*β* (Pearson *r* = 0.38, *p* < 1 × 10^−16^), and between the percent of sparse temporal coefficients and *α* (Pearson *r* = 0.37, *p* < 1 × 10^−16^). Moreover, we observe two likely basins of low and high temporal expression sparsity for a range of *β* and α, centered around (*β* ∊ [0.1,0.3], *α* ∊ [0.1,0.5]) and (*β* ∊ [1.25,2.0],*α* ∊ [1.25,2.0]). These results suggest that subgraphs may be specific to two different scales of temporal dynamics: local (higher percentage of sparse coefficients) or global (lower percentage of sparse coefficients). However, we do not observe similar basins in the relationship between the percent of sparse subgraph edge weights and *β* (Pearson *r* = 0.51, *p* < 1 × 10^−16^), nor between the percent of sparse subgraph edge weights and *α* (Pearson *r* = 0.51, *p* < 1 × 10^−16^). These results suggest a potential strategy for choosing parameters that is based on minimizing the residual error and maintaining a balance between spatial and temporal generalizability and specificity of the subgraphs. Therefore, we average the randomly sampled parameters associated with the lowest 25% residual error, greatest 25% sparse expression coefficients, and greatest 25% sparse edge weights. These choices resulted in a number of subgraphs *m* = 16.64 ± 0.07, a temporal sparsity of *β* = 1.383 ± 0.009, and a subgraph regularization of *α* = 1.368 ± 0.009.

### 3.2. Subgraphs stratify proximal and distributed functional interactions

An open question in network neuroscience is “What is the spatial organization of naturally formed subgraphs of dynamic brain networks?” We expect that subgraphs stratify coherent groups of functional interactions into clusters that reflect more local processing over shorter distances and more distributed processing over longer distances. To investigate the verity of this expectation, we compute the Euclidean distance between all pairs of 112 brain regions defined by the Harvard-Oxford anatomical atlas (Fig. 3A). Based on these distances, we measure the topographical sensitivity for each subgraph based on the correlation between a subgraph’s edge weights and the Euclidean distance between the brain regions represented by those edges. To compare topographical sensitivity across subgraphs of all subjects, we rank subgraphs in increasing order of their correlation with spatial distance and we analyze the distribution of correlations for each subgraph in comparison to a null model constructed from surrogate data (Fig. 3B). The surrogate data null model represents the null distribution of correlations when subgraphs lose their characteristic topology and instead exhibit random wiring.

We find that ten of the sixteen subgraphs exhibit a more negative correlation than expected by the surrogate model (Bonferroni corrected *t*-test; *p* < 0.05), demonstrating that stronger functional interactions are expressed over shorter distances. Five other subgraphs exhibit a more positive correlation than expected by the surrogate model (Bonferroni corrected *t*-test; *p* < 0.05), demonstrating that stronger functional interactions are expressed over longer distances. These results suggest that fifteen of the sixteenn subgraphs are composed of uniquely local or uniquely distributed functional interactions indicative of different scales of information processing (Fig. 3C). In light of recent literature [69, 61, 91], these diverse spatial scales of subgraphs may directly relate to the temporal scale of dynamics that the subgraphs support: that is, spatially proximal brain regions can fluctuate in their interaction at a different time-scale than spatially distant brain regions.

### 3.3. Module-based constraints on subgraph architecture

Next, we ask “What constraints might modular organization impart on subgraphs of functional brain networks? Are interactions specified by a subgraph restricted to brain regions of the same module, or can they span multiple modules?” By answering these questions, we can begin piecing together a mechanistic role for subgraphs in the architecture and dynamics of functional brain networks. Based on the perspective that modularity provides a substrate for both integrated and segregated modes of information processing [13, 109], we hypothesize that the strength of functional interactions will co-vary based on whether they are localized between brain regions within the same module or between brain regions in different modules (Fig. 4A). If such temporal co-variance were to exist within and between modules, functional subgraphs would reflect the various clusters of functional interactions that fluctuate over the different time-scales of information processing.

**Figure 4:**
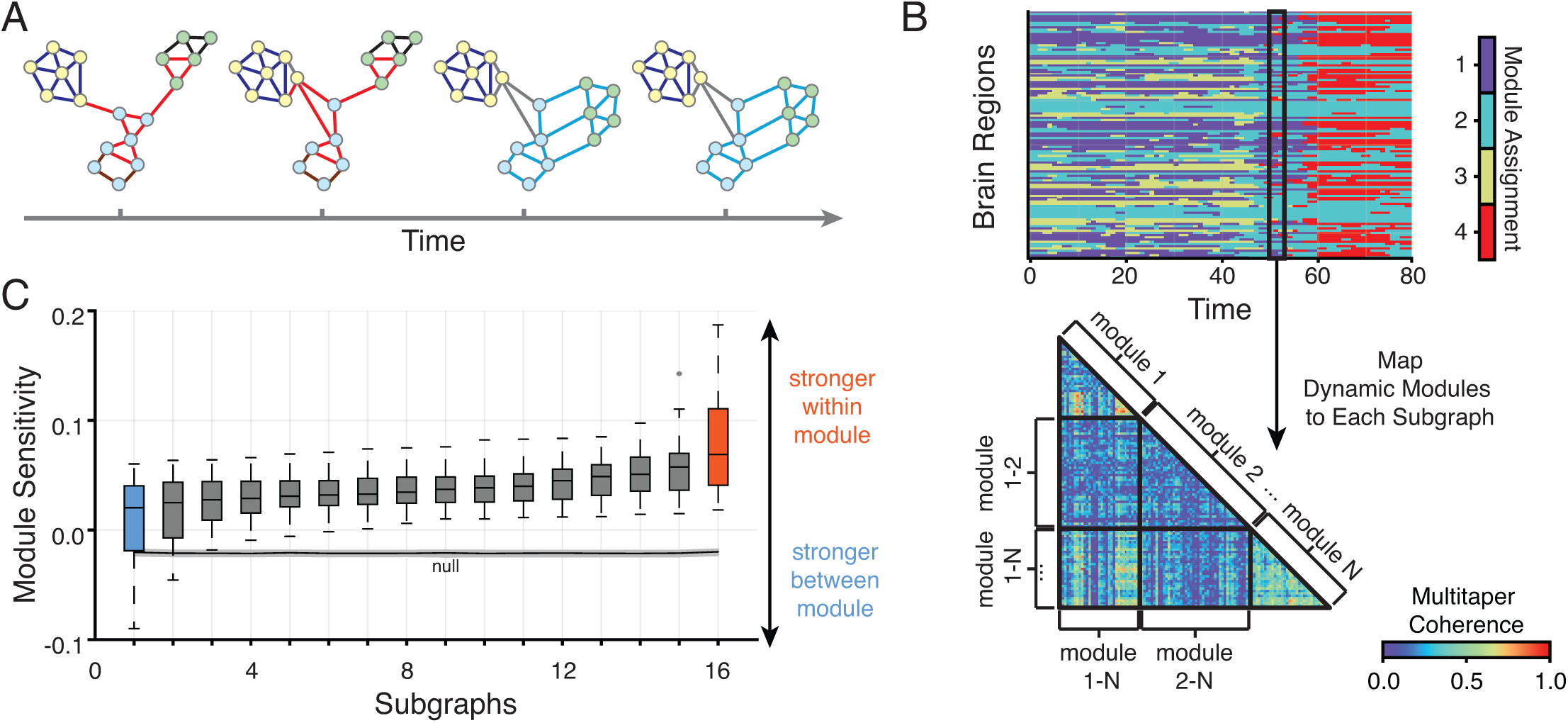
Network modules and network subgraphs. (*A*) A toy dynamic network that illustrates differential changes to the architecture of network modules and network subgraphs over time. Modules describe a hard-partition of strongly connected network nodes into different groups or “communities” (colored circles) whose composition may change over time. Subgraphs describe a soft-partition of temporally co-varying network edges into different groups or clusters (colored lines). Network edges of a single subgraph may occupy a single module (dark blue) or span multiple modules (light blue). (*B*) To quantify the degree to which a subgraph captures the modular organization of the network, we mapped module assignments from each time window (*left*) onto each subgraph and computed a time-varying module sensitivity index for each subgraph: positive values indicate greater within-module edge strength, while negative values indicate greater between-module edge strength. (*C*) Distribution of time-averaged module sensitivity index of subgraphs in increasing order over subjects (*N* = 20). Subgraphs exhibited significantly greater module sensitivity to true modular organization compared to a null model in which module assignments were permuted uniformly at random between nodes and between time windows (Bonferroni corrected *t*-tests; *p* < 0.05). However, subgraphs exhibited significantly different module sensitivities from one another (one-way ANOVA; *F*_19_ *=* 8.42, *p* < 1 × 10^−15^), suggesting that subgraphs with the greatest module sensitivity tend to capture strongest edges within modules (red) and subgraphs with the lowest module sensitivity tend to distribute strongest edges more evenly within and between modules (blue).

To quantify the sensitivity of functional subgraphs to topology within or between modules, we compute the time-varying module sensitivity index for each subgraph of each subject (see Methods). Intuitively, values closer to 1 imply that a subgraph expresses stronger functional interactions between brain regions assigned to the same module, and values closer to −1 imply that a subgraph expresses stronger functional interactions between brain regions assigned to different modules. Probing this index enables us to map subgraph topology to the modular architecture of the network at any given point in time (Fig. 4B). To compare the average module sensitivity across subgraphs of all subjects, we rank all of the subgraphs in increasing order of their average module sensitivity index over time, and we analyze the distribution of sensitivity indices for each subgraph in comparison to a null model built from surrogate data (Fig. 4C). The surrogate data null model represents the null distribution of module sensitivity indices when module assignments are permuted uniformly at random across brain regions.

We find that all subgraphs exhibit significantly greater average module sensitivity than expected in the surrogate model (Bonferroni corrected *t*-test; *p* < 0.05). Moreover, we find that subgraphs significantly stratify the observed distribution of average module sensitivity (one-way ANOVA; *F*_19_ = 8.42, *p* < 5 × 10^−16^). Together, these results suggest that modular architecture heterogenously constrains the functional interactions of subgraphs – some subgraphs express topology that resembles function-specific information processing within modules while other subgraphs express topology that resembles integrative processing across modules.

### 3.4. Module-based constraints on subgraph dynamics

Based on the strong relationship between modular architecture and functional subgraphs, we next ask whether dynamical changes in subgraph expression capture the module-based reorganization of brain networks. We expect that the expression dynamics of subgraphs with the greatest module sensitivity would exhibit a stronger relationship with module reorganization dynamics than subgraphs with the lowest module sensitivity. To investigate this hypothesis, we measure the energy and transience of subgraph expression and the flexibility of module reorganization. Subgraphs with the highest module sensitivity demonstrate a significant decrease in expression energy (Pearson *r* = −0.44, *p* < 0.05) and a significant increase in expression transience (Pearson *r* = 0.55, *p* < 0.05) with increasing module flexibility (Fig. 5A,B). In contrast, subgraphs with the lowest module sensitivity do not demonstrate significant relationships between expression energy and module flexibility (Pearson *r* = −0.16, *p* = 0.49), nor between expression transience and module flexibility (Pearson *r* = −0.13, *p* = 0.58). These results suggest that subgraphs with greater module sensitivity exhibit expression dynamics that more strongly relate to temporal changes in module organization. Specifically, subjects whose module-sensitive subgraphs have greater energy and lower transience, and thus high overall expression with less intermittent jumps in activity, tend to exhibit more fundamentally stable module architecture that reorganizes less frequently over time.

**Figure 5:**
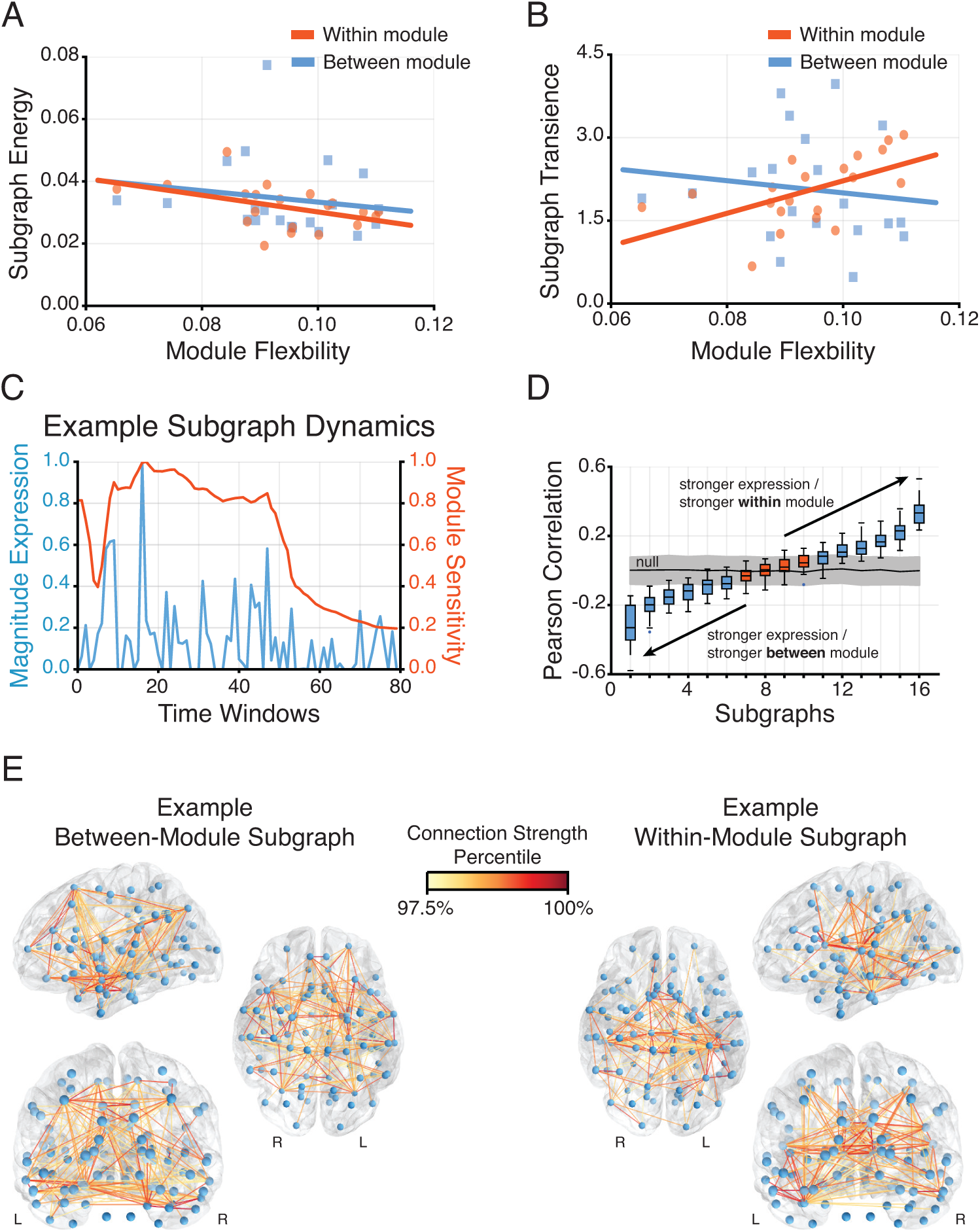
Dynamics of module sensitivity. (*A*) Relationship between module flexibility – the average rate nodes change their module allegiance – and subgraph expression energy – the magnitude a subgraph is expressed over a period of time – for subgraphs with high (red) and low (blue) average module sensitivity. Subgraphs with strongest edges within network modules significantly attenuate their expression in conjunction with more frequent changes in node allegiance to a module (Pearson *r* = −0.44, *p* < 0.05), compared to subgraphs with strong edges distributed between modules (Pearson *r* = −0.16, *p* = 0.49). (*B*) Relationship between module flexibility and subgraph expression transience – the distribution skew of expression coefficients within a window of time – for subgraphs with high (red) and low (blue) module sensitivity. Subgraphs with strongest edges within network modules are more transiently expressed in conjunction with more frequent changes in node allegiance to a module (Pearson *r* = 0.55, *p* < 0.05), compared to subgraphs with strongest edges distributed between modules (Pearson *r* = −0.13, *p* = 0.58). (*C*) Expression dynamics (blue) coincide with time-varying module sensitivity (red) in an example subgraph of a single subject, suggesting that a subgraph’s temporal expression may capture periods of heightened functional interaction within modules. To quantify whether the relationship between expression dynamics and module sensitivity generalizes to all subgraphs, we computed the Pearson correlation coefficient between subgraph expression and time-varying module sensitivity for each subgraph of each subject: positive values indicate increased sensitivity to topology within modules when expression is increased, while negative values indicate increased sensitivity to topology between modules when expression is increased. (*D*) Distribution of Pearson correlation coefficient values between subgraph expression and time-varying module sensitivity of subgraphs in increasing order over subjects (*N* = 20). Compared to a null model in which temporal expression weights were permuted uniformly at random, subgraphs 1-6 (blue) exhibited increased sensitivity to between-module topology with increased expression, subgraphs 11-16 exhibited increased sensitivity to within-module topology with increased expression, and subgraphs 7-10 exhibited non-significant relationships between module sensitivity and expression (Bonferroni corrected *t*-tests; *p* < 0.05). These results support the hypothesis that different subgraph topologies capture varying degrees of edge co-variance within or between network modules specific to time periods of heightened expression. (*E*) An example of a between-module subgraph with significant negative Pearson correlation values (*left*), and an example of a within-module subgraph with significant positive Pearson correlation values (*right*). The between-module subgraph exemplifies strong edges distributed between modules and the within-module subgraph exemplifies strong edges localized within modules.

Based on our finding that subgraphs reflect functional interactions within and between modules, it is natural to ask whether temporal expression of subgraphs captures network states that resemble inter-*versus* intra-module processes. We expect that subgraph expression dynamics differentially explain temporal fluctuations in the module sensitivity of subgraphs (Fig. 5C). To address this hypothesis, we measure the temporal similarity between expression dynamics and module sensitivity index for each subgraph using a Pearson correlation: values closer to 1 imply that increased expression of a subgraph is associated with increased sensitivity to functional interactions within modules. In contrast, values closer to −1 imply that increased expression of a subgraph is associated with increased sensitivity to functional interactions between modules. To compare temporal similarity across subgraphs of all subjects, we rank subgraphs in increasing order of their Pearson correlation and analyze the distribution of correlations for each subgraph in comparison to a null model built from surrogate data (Fig. 5D). The surrogate data null model represents the null distribution of correlations when subgraph expression dynamics lose their temporal structure through random permutation of the expression coefficients.

We find that six of the sixteen subgraphs exhibit a more negative correlation than expected by the surrogate model (Bonferroni corrected *t*-test; *p* < 0.05), indicating that increased expression implies greater sensitivity to between-module interactions. A separate six subgraphs exhibit a more positive correlation than expected by the surrogate model (Bonferroni corrected *t*-test; *p* < 0.05), indicating that increased expression implies greater sensitivity to within module interactions. Collectively, these results suggest a division of subgraph types into two groups characterized by increased inter- or intra-module interaction that scales with the magnitude of temporal expression (Fig. 5E). Fluctuations in subgraph expression amongst these two different groups form a potential mechanism for the temporal dynamics of information integration and segregation that is commonly observed within and between modules [125,77,16].

Overall, our findings suggest that the modular architecture of brain networks places fundamental constraints on the various modes of functional interactions between brain regions, as represented by subgraphs. Thus, network modules and network subgraphs may play complimentary roles in guiding dynamics of functional brain networks; network modules prescribe the meso-scale organization of functionally cohesive brain regions, network subgraphs pinpoint the modes in which these brain regions interact with finer granularity. Intriguingly, the dynamics of these individual subgraphs explain fluctuations in integrated and segregated module-level interactions, as well as inter-subject variability in module reorganization.

## 4. Discussion

In this study, we put forth a framework for uncovering topological modes of functional interactions from time-evolving brain networks that is based on an unsupervised machine learning tool called non-negative matrix factorization. We demonstrate that functional brain networks decompose into constituent parts or additive subgraphs that are differentially expressed over time. To effectively identify these subgraphs, we demonstrate an ability to control the number of subgraphs, the temporal sparseness of their expression, and the topological sparseness of their edge weights by manipulating three important parameters of the NMF optimization problem. Using the most robust parameter set for the decomposition, we investigate the topographical and topological basis of the recovered functional subgraphs. We find that subgraphs naturally stratify functional interactions between spatially proximal brain regions and spatially distributed brain regions. Upon closer examination, we observe that these clusters of coherent functional interactions underlie a brain architecture that supports integration and segregation of information processing across network modules. Moreover, subgraphs exhibit expression dynamics that reliably signal the modular reorganization of the network.

### 4.1. Machine learning to partition functional brain networks

Network neuroscientists are eager to use computational tools to objectively partition brain networks into naturally organized sub-regions that can deepen our understanding of form and function [23]. Conventional approaches to localize network sub-regions or components are based on graph theoretic tools, such as the minimum-cut procedure [97], that incrementally and continually divide a network along its edges until a chosen criteria is satisfied. While such algorithms can be particularly effective for examining clusters of brain regions in anatomically-based structural networks that remain relatively static over short periods of time [7, 55], they are not designed to partition dynamic functional networks whose architecture evolves over time. Dynamic community detection methods fill this void by enabling network scientists to identify modules – cohesive groups of highly interacting brain regions – whose composition could change over time [8].

Importantly, traditional methods for community detection pursue a *hard* partitioning of brain networks that unambiguously assigns brain regions to a single module based purely on the strength, rather than the topological arrangement, of functional interactions [92, 45]. However, recently it has been argued that brain regions may not necessarily have disjoint organization where they fulfill a single functional role. Rather, they may participate in many brain systems along different phases of behavioral and cognitive processing [124, 90, 41] – requiring models that are capable of *soft* partitioning the network such that brain regions are allowed to participate in different systems to varying degree [4, 1]. In the brain, such soft partitions can come in the form of so-called link communities [37] or hyper-graphs [11, 35, 36], although neither of these approaches addresses the existence of dynamics in the partition.

To accommodate greater model flexibility in characterizing network organization and dynamics, we turned to soft partitioning approaches – such as NMF – that afford the ability to statistically learn important rules regarding the behavior of a system through observational data. Unlike hard partitioning approaches – such as community detection – that may be tailored to understand different facets of network organization under particular constraints, soft partitioning deconstructs the statistical space of dynamic networks, provided there is working knowledge of the underlying distribution of that space. Put simply, hard partitioning approaches ask “Are brain networks organized in a particular way?”, while soft partitioning approaches ask “What are the rules underlying the observed network organization?” Thus, community detection clusters brain regions into modules based on the strength of their interactions, while NMF clusters functional interactions into subgraphs based on their cohesive fluctuations over time.

### 4.2. Spatial and temporal constraints on functional interactions

Fundamentally, subgraphs represent constraints on the functional interactions that the observed network is able to support. Intriguingly, prior work demonstrates that functional subgraphs share resounding similarity to empirically-defined brain systems [27] – such as the executive system [108, 105, 88] and default mode network [95, 96] – supporting the theory that subgraphs prescribe a foundational substrate for the functional interactions underlying cognitive processes and resulting behavior. In our study, we observe that functional subgraphs support heterogeneous topography of interactions over short and long distances. The observed stratification of distance-based interactions over several subgraphs suggests that these subgraphs also represent unique topological relationships. Our findings are supported by previous studies that use both descriptive statistics [75, 7, 67] and generative models [122, 17] to investigate the relative importance of topography and topology in anatomical networks. Functionally, subgraphs that separately support short-range and long-range interactions could be further explored in their potential role in shaping human intelligence, which has been linked to long-distance interactions that promote global efficiency [74, 119, 104].

A long purported role of multi-scale network topology is to support segregated processing of functionally specialized information and integrated processing of distinct pieces of information [23, 18]. Prior studies demonstrate an ability to capture the functional substrates of integration and segregation during learning [9, 13, 48], linguistic processing [29, 42], aging [31, 19, 81], executive function [22, 20], attention [77, 115], and rest [16] using modular decomposition. While modular organization helps compartmentalize function-specific computations within individual modules, the finer-scale topology that actuates this computation has remained elusive. Our results demonstrate that subgraph topologies differentially express functional interactions within and between modules. These within- and between-module subgraphs may help define a clearer functional role for putative provincial and connector hubs in the context of local and distributed computations [16]. In addition to a topological basis for module-based processing, our results suggest that subgraph dynamics may distinguish different stages of integrated and segregated processing. The ability to track the relative expression of subgraphs during these stages is crucial for understanding the temporal dynamics that embody so called *cognitive control* processes [82] that govern our ability to modulate attention [68, 47, 34] and switch between tasks [102,64, 121].

Network flexibility, the ability for modular architecture to reorganize, has recently been considered a putative functional driver of cognitive control [33, 13, 22], and is demonstrably altered in psychiatric disorders such as schizophrenia [20], which are also characterized by deficits in executive function [108, 6]. In this study, we explore possible relationships between modular reorganization and subgraph dynamics and find that subgraphs that are more sensitive to topology within modules have stronger relationships between their temporal pattern of expression and network flexibility. First, these subgraphs are more energetically expressed in individuals who exhibit less flexible functional modules, potentially implicating expression energy as a stabilizer of network architecture. Such a theory is corroborated by prior work that finds subgraphs of the executive system are more energetically expressed in young adults compared to children [27], which might be explained by less impulsive and more proactive behavior [117, 32]. Second, individuals with module-sensitive subgraphs that are more transiently expressed tend to exhibit more flexible functional modules, instantiating expression transience as an intermittent modulator of network architecture. Subgraphs with more transient dynamics might signal important transition states that potentially drive changes in overall network topology in an inherently rough energy landscape [3, 123, 54, 98].

### 4.3. Conclusions

In this study, we introduce a machine learning approach for decomposing dynamic functional networks into subgraphs and characterizing their architecture in the context of topography and topology. We show that subgraphs stratify groups of functional interactions that are expressed over a variety of spatial distances. Furthermore, subgraphs express topologies that obey the modular architecture of functional brain networks. These topological constraints lend subgraphs an important ability to signal dynamical states of inter- and intra-module processing. In general, our framework can be used to examine the functional substrates underlying various behavioral and cognitive states, such as the coordination of different groups of functional interactions during a task. Importantly, the NMF-based approach would highlight which brain regions are engaged during these states and how these engagements shift over time. Further methodological development could highlight sequences of subgraph expression that could be used to understand dynamics associated with cognitively effortful tasks, and their alteration in patients with neurological disorders or psychiatric disease, and pinpoint causal drivers of network dynamics [76, 50].

## 4.4. Acknowledgments

A.N.K, M.G.M, and DSB would like to acknowledge support from the John D. and Catherine T. MacArthur Foundation, the Alfred P. Sloan Foundation, the Army Research Laboratory and the Army Research Office through contract numbers W911NF-10-2-0022 and W911NF-14-1-0679, the National Institute of Health (2-R01-DC-009209-11, 1R01HD086888-01, R01-MH107235, R01-MH107703, R01MH109520, 1R01NS099348 and R21-M MH-106799), the Office of Naval Research, and the National Science Foundation (BCS-1441502, CAREER PHY-1554488, BCS-1631550, and CNS-1626008).The content is solely the responsibility of the authors and does not necessarily represent the official views of any of the funding agencies.

